# Evaluating the Synergetic Effect of *Nigella sativa* Extract and Metal Oxide Nanoparticles Against Different Microbes

**DOI:** 10.1101/2025.11.28.691110

**Authors:** Muhammad Owais Jan

**Affiliations:** Institute of Integrative Biosciences, Faculty of Life Sciences, CECOS University, Peshawar, Pakistan

**Keywords:** *Nigella sativa*, zinc oxide nanoparticles, antimicrobial activity, synergetic effect, metal oxide nanoparticles

## Abstract

**Background:** The emergence of antibiotic-resistant bacteria necessitates the exploration of alternative antimicrobial agents. *Nigella sativa* (black seed) and metal oxide nanoparticles have independently shown antimicrobial properties, but their synergetic effects remain understudied.

**Objective:** This study investigated the individual and combined antimicrobial activities of *Nigella sativa* extract with zinc oxide (ZnO), manganese-doped zinc oxide (Mn-ZnO), and cobalt-doped zinc oxide (Co-ZnO) nanoparticles against *Escherichia coli*, *Staphylococcus aureus*, and *Pseudomonas aeruginosa*.

**Methods:** Nanoparticles were synthesized via co-precipitation method and characterized using UV-Vis spectroscopy, X-ray diffraction (XRD), transmission electron microscopy (TEM), and scanning electron microscopy (SEM). *N. sativa* extract was prepared through maceration in 70% methanol. Antimicrobial activity was assessed using agar well diffusion assay and minimum inhibitory concentration (MIC) determination.

**Results:** Nanoparticles demonstrated significant antimicrobial activity against *E. coli* (inhibition zones: 2.5-2.7 cm) but showed no activity against *S. aureus* and *P. aeruginosa*. Conversely, *N. sativa* extract exhibited antimicrobial activity against *S. aureus* and *P. aeruginosa* (inhibition zones: 2.0-2.2 cm) but not against *E. coli*. No synergetic effect was observed when nanoparticles and *N. sativa* extract were combined.

**Conclusion:** The study revealed complementary but non-synergistic antimicrobial activities between *N. sativa* extract and metal oxide nanoparticles. Future research should investigate the molecular mechanisms preventing synergy and explore potential antagonistic interactions.

## 1. Introduction

Nanoscience represents one of the most rapidly developing multidisciplinary fields due to its extensive theoretical and practical applications. The synthesis of nanoparticles through physical, chemical, biological, and hybrid approaches has opened new avenues for antimicrobial research, particularly in addressing the growing challenge of antibiotic resistance.

*Nigella sativa* L., commonly known as black seed or black cumin, belongs to the Ranunculaceae family and has been utilized in traditional medicine across Middle Eastern and Asian cultures for centuries. The seeds contain numerous bioactive compounds, with thymoquinone being the most pharmacologically active constituent, demonstrating a broad spectrum of therapeutic properties including antimicrobial, anti-inflammatory, antioxidant, and anticancer activities.

Metal oxide nanoparticles, particularly zinc oxide (ZnO) and its doped variants, have gained considerable attention due to their antimicrobial properties. ZnO is a versatile semiconducting material with a band gap of 3.37 eV, exhibiting excellent chemical and thermal stability, biocompatibility, and antimicrobial activity. Doping ZnO with transition metals such as manganese (Mn) and cobalt (Co) can enhance surface area, modify optical properties, and improve antimicrobial efficacy.

Despite independent studies demonstrating the antimicrobial potential of both *N. sativa* and metal oxide nanoparticles, their combined effects remain largely unexplored. This study aimed to evaluate the individual and synergetic antimicrobial activities of *N. sativa* extract combined with ZnO, Mn-doped ZnO, and Co-doped ZnO nanoparticles against three clinically relevant bacterial strains: *Escherichia coli*, *Staphylococcus aureus*, and *Pseudomonas aeruginosa*.

## 2. Materials and Methods

### 2.1 Materials

#### 2.1.1 Chemicals and Reagents

Methanol (analytical grade), zinc nitrate hexahydrate, cobalt nitrate hexahydrate, manganese nitrate tetrahydrate, potassium hydroxide (KOH), polyethylene glycol (PEG), Luria broth (LB), agar, barium chloride dihydrate, sulfuric acid (H₂SO₄), and ethanol were obtained from commercial suppliers.

#### 2.1.2 Bacterial Strains

*Escherichia coli*, *Staphylococcus aureus*, and *Pseudomonas aeruginosa* clinical isolates were obtained from the laboratory collection.

#### 2.1.3 Plant Material

*Nigella sativa* seeds were purchased from a local herbal shop in Peshawar, Pakistan, and authenticated.

### 2.2 Preparation of *Nigella sativa* Extract

Seeds were washed thoroughly under running water, rinsed twice with distilled water, and dried in an oven at 40°C for 24 hours. Dried seeds were ground into fine powder using an electric grinder. The powdered seeds

(100 g) were macerated in 70% methanol (500 mL) for 24 hours at room temperature with occasional stirring. The mixture was filtered through Whatman No. 1 filter paper, and the filtrate was concentrated using a rotary vacuum evaporator. The crude extract was dissolved in dimethyl sulfoxide (DMSO) to prepare stock concentrations of 1, 2, and 3 mg/mL.

### 2.3 Synthesis of Nanoparticles

#### 2.3.1 Zinc Oxide Nanoparticles (ZnO NPs)

Zinc nitrate hexahydrate (1 M) was dissolved in 50 mL distilled water. Potassium hydroxide (2 M) dissolved in 50 mL distilled water was added dropwise to the zinc nitrate solution under constant stirring for 2 hours. The precipitate was collected by centrifugation at 10,000 rpm for 15 minutes, washed three times (once with water, twice with ethanol, once with water), and dried at 60°C for 24 hours. The dried powder was calcined at 200°C for 16 hours.

#### 2.3.2 Manganese-Doped Zinc Oxide Nanoparticles (Mn-ZnO NPs)

Zinc nitrate hexahydrate (1 M) and manganese nitrate tetrahydrate (2 M) were dissolved in 50 mL distilled water. Potassium hydroxide solution (4 M in 50 mL water) was added dropwise under stirring for 2 hours. The precipitate was processed as described above. After drying, nanoparticles were coated with PEG by dissolving 10× weight of PEG in 100 mL water and stirring for 24 hours. The coated nanoparticles were dried and calcined at 200°C for 16 hours.

#### 2.3.3 Cobalt-Doped Zinc Oxide Nanoparticles (Co-ZnO NPs)

Zinc nitrate hexahydrate (1 M) was dissolved in 25 mL water, and 10% cobalt nitrate hexahydrate was added. KOH solution (1 M in 50 mL water) was added dropwise under stirring for 2 hours. The precipitate was washed, dried, and calcined as described for ZnO NPs.

### 2.4 Characterization of Nanoparticles

#### 2.4.1 UV-Visible Spectroscopy

Nanoparticle formation was confirmed using UV-Vis spectrophotometer (wavelength range: 200-800 nm).

#### 2.4.2 X-ray Diffraction (XRD)

Crystal structure was analyzed using a JEOL (JDX-3532) X-Ray diffractometer with Cu-Kα radiation at tube voltages of 20-40 kV and 2.5-30 mA, scanning from 0° to 160°.

#### 2.4.3 Transmission Electron Microscopy (TEM)

Particle size and morphology were examined using JEOL-2100 TEM operating at 100-200 keV. Samples were prepared by depositing diluted nanoparticle suspension on copper grids and air-drying.

#### 2.4.4 Scanning Electron Microscopy (SEM)

Surface morphology was analyzed using JSM-5910 SEM at magnifications of 5,000× to 60,000×.

### 2.5 Antimicrobial Activity Assessment

#### 2.5.1 Bacterial Culture Preparation

Bacterial strains were cultured in LB broth at 37°C overnight. Bacterial suspensions were adjusted to 0.5 McFarland standard (approximately 1.5 × 10⁸ CFU/mL) using normal saline.

#### 2.5.2 Agar Well Diffusion Assay

Pure bacterial colonies were isolated using streak plate method. Bacterial suspensions were spread uniformly on LB agar plates using sterile cotton swabs. Wells (6 mm diameter) were created using a sterile cork borer. Test samples (100 μL) including nanoparticles (10 mg/mL), *N. sativa* extract (3 mg/mL), combinations, and controls (ethanol) were added to respective wells. Plates were incubated at 37°C for 24 hours. Zones of inhibition were measured in millimeters.

#### 2.5.3 Minimum Inhibitory Concentration (MIC)

Bacterial suspensions (100 μL) were added to wells of a 96-well microtiter plate containing LB broth. Test samples (200 μL) were added to designated wells in triplicate. The plate layout included: bacteria alone (negative control), bacteria + ethanol (positive control), bacteria + individual nanoparticles, bacteria + *N. sativa* extract, and bacteria + combined treatments. Plates were incubated at 37°C for 24 hours with shaking. Optical density was measured at 600 nm initially and after 24 hours using a microplate reader.

### 2.6 Statistical Analysis

All experiments were performed in triplicate. Data are presented as mean values. Inhibition zones were measured and recorded in centimeters.

## 3. Results

### 3.1 Nanoparticle Characterization

#### 3.1.1 UV-Visible Spectroscopy

UV-Vis spectroscopy confirmed successful nanoparticle synthesis with characteristic absorption peaks: ZnO NPs at 370 nm, Mn-ZnO NPs at 373 nm, and Co-ZnO NPs at 368 nm. These peaks correspond to the band gap of zinc oxide nanostructures.

#### 3.1.2 X-ray Diffraction Analysis

XRD analysis of Co-ZnO NPs revealed strong, sharp peaks indicating high crystallinity and polycrystalline structure. Diffraction peaks were observed at 2θ = 25.2°, 27.5°, 32.3°, 35.5°, 37.7°, 42°, 47°, 62°, 65°, and 68.0°, corresponding to the hexagonal wurtzite structure of ZnO. Peaks at (100), (002), and (101) planes confirmed pure wurtzite phase formation with no secondary phase impurities.

#### 3.1.3 Transmission Electron Microscopy

TEM images of Co-ZnO NPs showed oval-shaped particles with average size ranging from 40-60 nm, confirming nanoscale dimensions and relatively uniform morphology.

#### 3.1.4 Scanning Electron Microscopy

SEM analysis revealed Co-ZnO NPs with nearly spherical morphology and particle sizes between 200-300 nm. The agglomeration observed in SEM compared to TEM measurements suggests particle clustering during sample preparation.

### 3.2 Antimicrobial Activity

#### 3.2.1 Agar Well Diffusion Assay Results Against *Escherichia coli*

& ZnO NPs: 2.5 cm inhibition zone

& Mn-ZnO NPs: 2.6 cm inhibition zone

& Co-ZnO NPs: 2.7 cm inhibition zone

& *N. sativa* extract: 1.3-1.8 cm inhibition zone

& Combined treatments: 2.6 cm inhibition zone (similar to nanoparticles alone)

& Ethanol control: 3.0 cm inhibition zone

**Against *Staphylococcus aureus*:**

& All nanoparticles alone: No inhibition zone

& *N. sativa* extract: 2.0 cm inhibition zone

& Combined treatments: 2.1-2.2 cm inhibition zone (similar to *N. sativa* alone)

**Against *Pseudomonas aeruginosa*:**

& All nanoparticles alone: No inhibition zone

& *N. sativa* extract: 2.2 cm inhibition zone

& Combined treatments: No synergetic enhancement observed

#### 3.2.2 Minimum Inhibitory Concentration Results

**For *E. coli*:** Ethanol showed complete growth inhibition in wells 1 and 2 (100 μL concentration), with partial growth in well 3, establishing the MIC at approximately 100 μL. ZnO NPs exhibited similar efficacy. *N. sativa* extract and combined treatments showed bacterial growth in all wells, indicating insufficient antimicrobial activity at tested concentrations.

**For *S. aureus*:** Only ethanol demonstrated growth inhibition in initial wells. All nanoparticles, *N. sativa* extract alone, and combined treatments showed visible bacterial growth, with optical density increases from initial to final readings.

**For *P. aeruginosa*:** Results were consistent with *S. aureus*, where ethanol was the only effective treatment at lower concentrations. Nanoparticles showed no MIC effect, while *N. sativa* extract showed moderate activity but no synergetic enhancement when combined.

## 4. Discussion

This study investigated the individual and synergetic antimicrobial activities of *Nigella sativa* extract and metal oxide nanoparticles against three clinically relevant bacterial pathogens. The results revealed distinct, non-overlapping antimicrobial spectra for the two treatment modalities, with no synergetic enhancement observed upon combination.

### 4.1 Antimicrobial Activity of Nanoparticles

All synthesized nanoparticles (ZnO, Mn-ZnO, and Co-ZnO) demonstrated significant antimicrobial activity against *E. coli* with inhibition zones ranging from 2.5-2.7 cm. This activity can be attributed to multiple mechanisms including: (1) generation of reactive oxygen species (ROS) that damage bacterial cell membranes, (2) release of zinc ions that disrupt cellular metabolism, and (3) direct physical interaction with bacterial cell walls leading to membrane disruption.

The Gram-negative cell wall structure of *E. coli*, characterized by a thin peptidoglycan layer and an outer lipopolysaccharide membrane, may be more susceptible to nanoparticle penetration and ROS-mediated damage. Previous studies have demonstrated that ZnO nanoparticles accumulate in bacterial cell membranes, causing structural damage and ultimately cell death.

Surprisingly, the nanoparticles showed no antimicrobial activity against *S. aureus* (Gram-positive) and *P. aeruginosa* (Gram-negative). This selective activity contradicts some literature suggesting greater susceptibility of Gram-positive bacteria to ZnO nanoparticles. The thick peptidoglycan layer of *S. aureus* and the robust biofilm-forming capability of *P. aeruginosa* may provide protective barriers against nanoparticle action. Additionally, strain-specific resistance mechanisms or differences in experimental conditions may account for these observations.

### 4.2 Antimicrobial Activity of *Nigella sativa* Extract

The methanolic extract of *N. sativa* demonstrated antimicrobial activity against *S. aureus* and *P. aeruginosa* but showed minimal effect against *E. coli*. This pattern is consistent with the presence of bioactive compounds, particularly thymoquinone, which has been reported to exhibit antibacterial properties through multiple mechanisms including membrane disruption, inhibition of cellular enzymes, and interference with quorum sensing.

The effectiveness against *S. aureus* aligns with traditional uses of *N. sativa* in treating bacterial infections. Thymoquinone and other phenolic compounds present in the extract may interact more effectively with the Gram-positive cell wall structure. The activity against *P. aeruginosa* is particularly significant given this organism’s notorious antibiotic resistance.

The lack of activity against *E. coli* suggests that this bacterial species may possess specific resistance mechanisms or metabolic pathways that neutralize the active compounds in *N. sativa* extract. Alternatively, the extraction method may not have captured all antimicrobial components effective against *E. coli*.

### 4.3 Absence of Synergetic Effect

The most significant finding of this study is the complete absence of synergetic antimicrobial activity when *N. sativa* extract was combined with nanoparticles. In all cases, the combined treatment showed activity equivalent to the more effective individual component, suggesting neither synergy nor antagonism, but rather independent action.

Several hypotheses may explain this observation:

1. **Competing Mechanisms:** The extract components may interfere with nanoparticle surface interactions or ROS generation, preventing the nanoparticles from effectively penetrating bacterial cells.
2. **Chemical Interactions:** Organic compounds in the extract, particularly phenolic compounds and proteins, may adsorb onto nanoparticle surfaces, creating a corona that shields the nanoparticles and reduces their antimicrobial activity.
3. **Complementary but Non-Synergistic Activities:** Each treatment may target the same cellular components through different mechanisms, creating a ceiling effect where no additional benefit occurs from combination.
4. **Extract-Mediated Scavenging:** Antioxidant compounds in *N. sativa* extract may neutralize ROS generated by nanoparticles, effectively canceling out this key antimicrobial mechanism.

### 4.4 Methodological Considerations

The MIC results revealed that ethanol (used as a solvent for *N. sativa* extract) exhibited antimicrobial activity against all tested bacteria. This raises important considerations about the contribution of the solvent to observed activities. However, the clear differences in activity patterns between pure ethanol and *N. sativa* extract-containing ethanol suggest that the extract itself possesses distinct antimicrobial properties.

The concentration-dependent nature of antimicrobial activity was evident in MIC assays, where 100 μL represented the threshold for effective growth inhibition by ethanol and ZnO NPs against *E. coli*. Higher concentrations or alternative delivery methods may be necessary to achieve broader-spectrum activity.

### 4.5 Clinical and Research Implications

The complementary antimicrobial spectra of *N. sativa* extract and metal oxide nanoparticles suggest potential for combined therapeutic applications, despite the absence of synergy. A treatment regimen incorporating both modalities could potentially address infections involving multiple bacterial species or provide broader coverage against unknown pathogens.

The absence of synergy highlights the importance of understanding molecular interactions in combination therapies. Future research should focus on:

1. Identifying specific compounds in *N. sativa* extract that may interfere with nanoparticle activity
2. Investigating sequential rather than simultaneous application of treatments
3. Exploring alternative nanoparticle coatings or modifications that prevent extract interference
4. Examining the role of antioxidant compounds in mediating antimicrobial activities

### 4.6 Study Limitations

Several limitations should be acknowledged. First, only three bacterial species were tested; broader screening against additional Gram-positive and Gram-negative bacteria would provide more comprehensive insights.

Second, the study used crude *N. sativa* extract rather than purified compounds, making it difficult to identify specific molecular interactions. Third, the mechanisms underlying the observed antimicrobial activities were not directly investigated through biochemical or molecular assays.

## 5. Conclusion

This study demonstrated that *Nigella sativa* extract and metal oxide nanoparticles possess complementary but non-synergistic antimicrobial activities. Nanoparticles effectively inhibited *E. coli* growth, while *N. sativa* extract showed activity against *S. aureus* and *P. aeruginosa*. No synergetic enhancement was observed when treatments were combined, suggesting potential molecular interactions that prevent cooperative antimicrobial action.

These findings have important implications for the development of combination antimicrobial therapies. While the complementary activity spectra could be leveraged for broad-spectrum applications, the lack of synergy indicates that simple combination may not enhance efficacy beyond individual treatments. Future research should focus on elucidating the molecular mechanisms preventing synergy and exploring alternative formulation strategies such as sequential application, chemical modification of nanoparticles, or use of purified bioactive compounds from *N. sativa*.

Understanding the interactions between natural products and nanomaterials is crucial for advancing antimicrobial therapeutic development, particularly in the context of rising antibiotic resistance.

## Acknowledgments

The author gratefully acknowledges Mr. Aamir Wahab (supervisor), Dr. Zia Ullah, and Dr. Kashif for their technical guidance and support throughout this research. Sincere thanks to the Department of Biotechnology, Institute of Integrative Biosciences, CECOS University, Peshawar, for providing laboratory facilities and resources.

---

**Figure 3.1.**
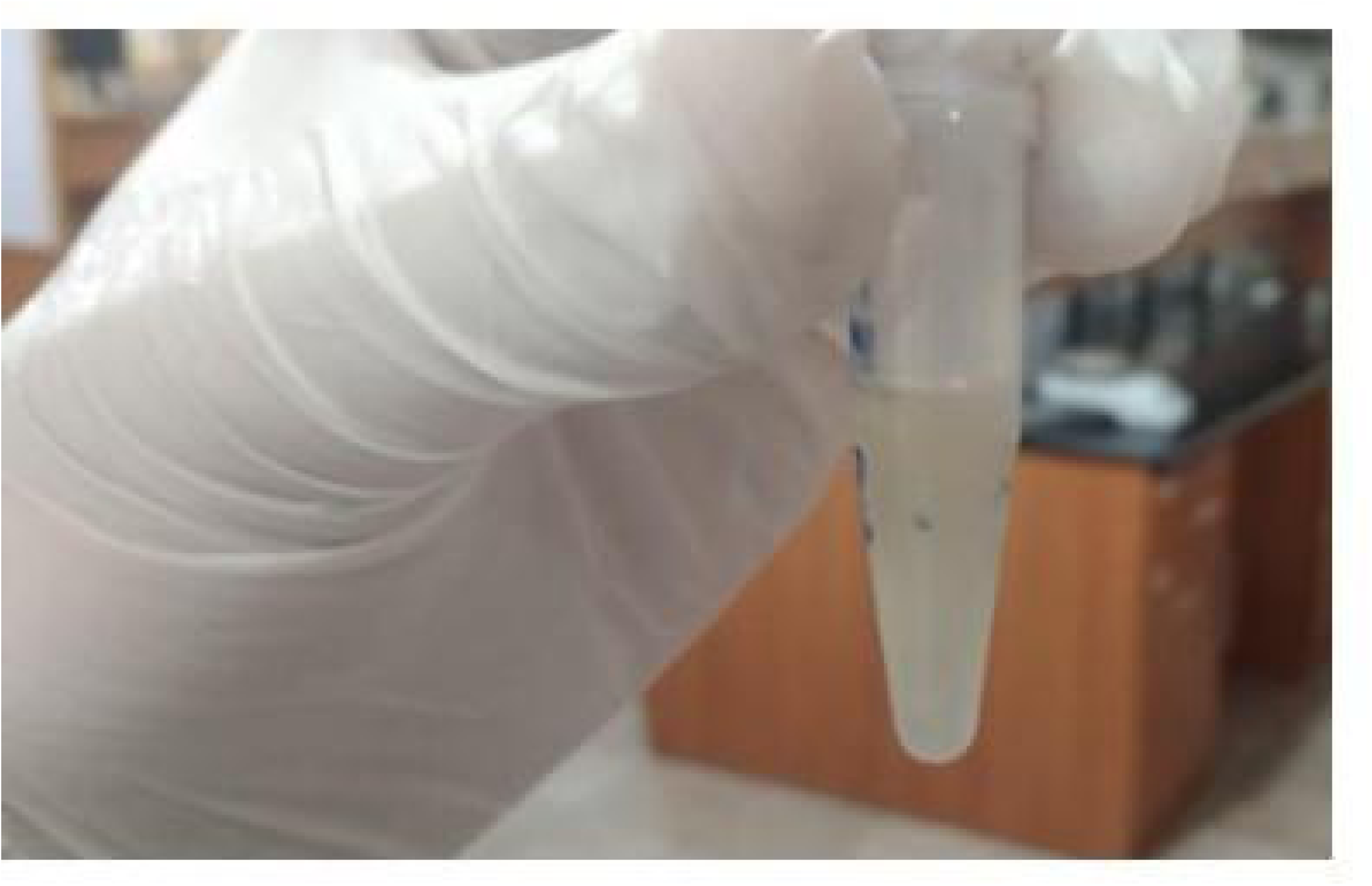
Macerated *Nigella sativa* and DMSO

**Figure 3.2.**
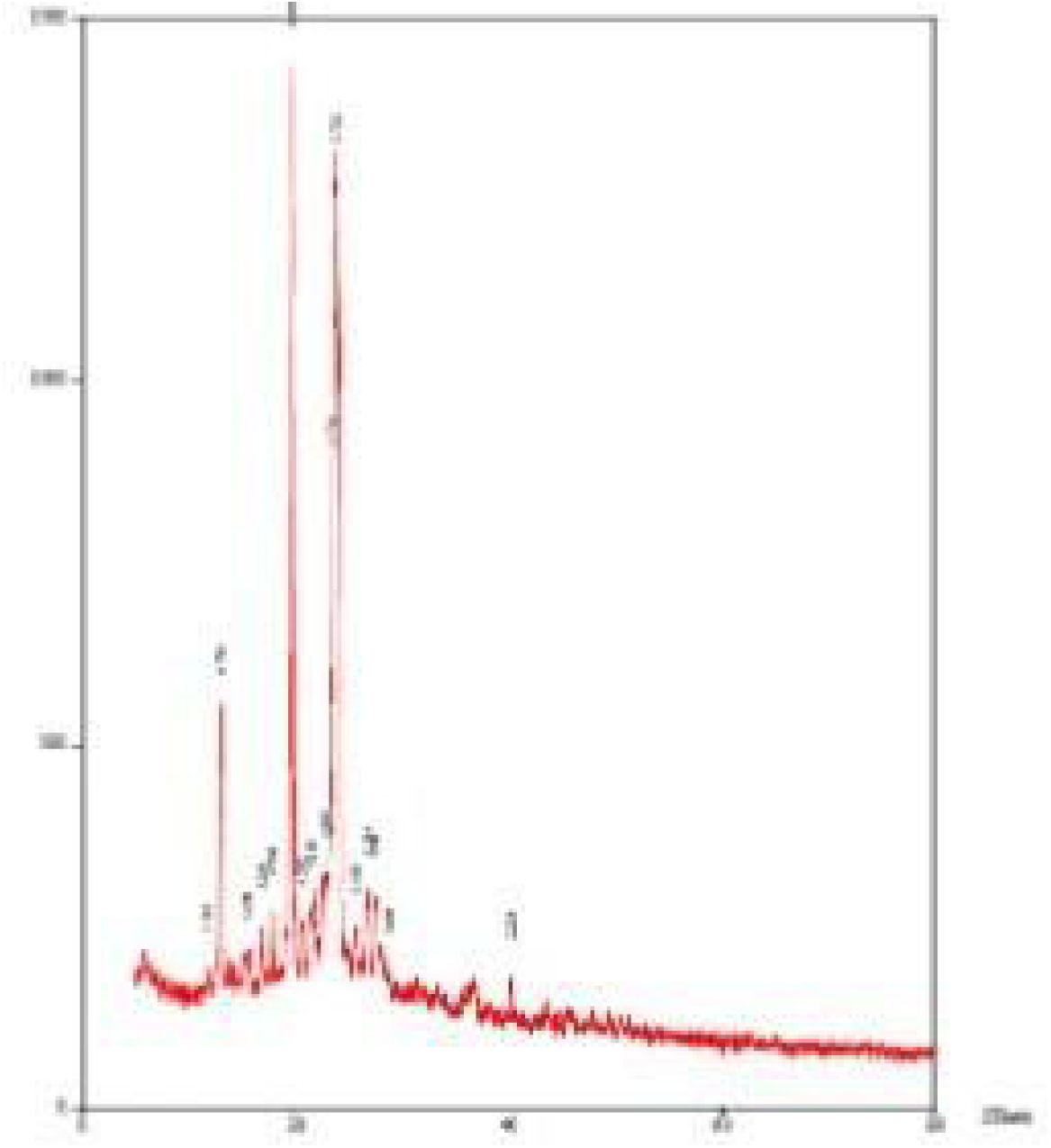
Marngoo.ese doped zinc oxide chMacterization through J<RD,

**Figure 3.3.**
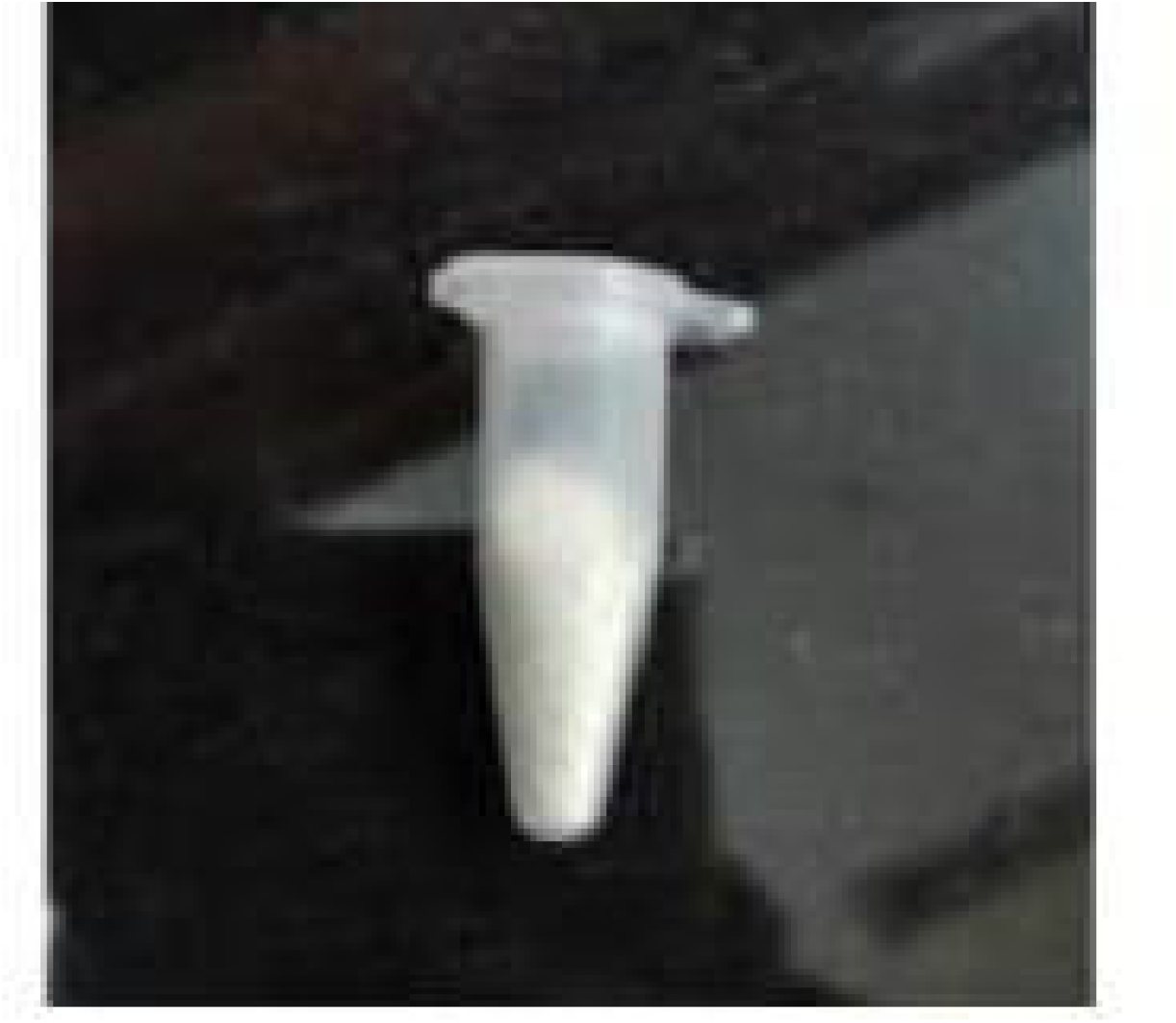
Zinc nanoparticles

**Figure 3.4.**
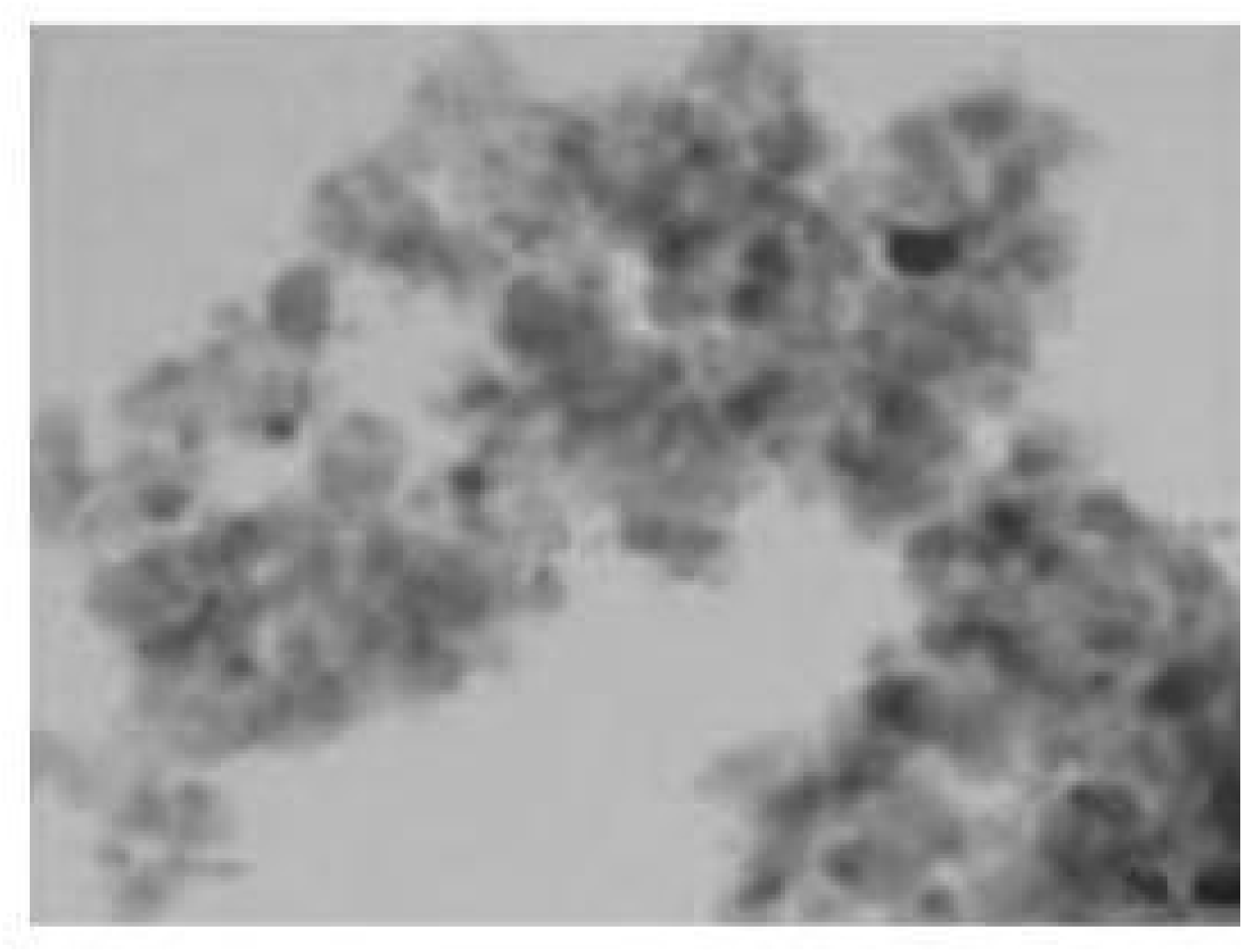
SEM analysis of zine oxide nanoparticles

**Figure 3.6.**
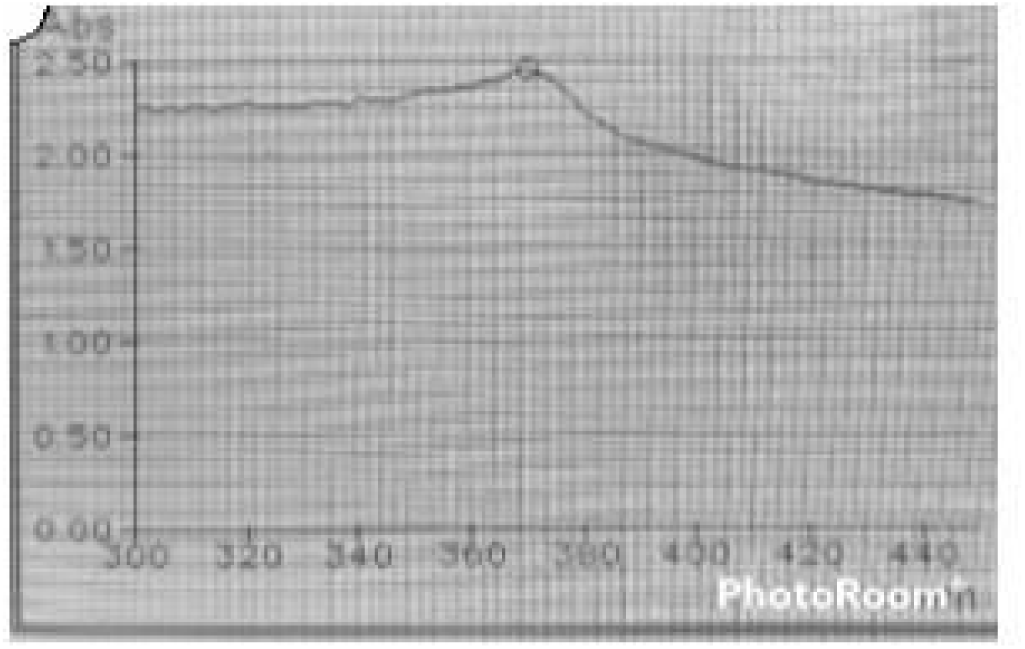
UV/Vis Spectroscopy results.of Zinc Oxide nanopartides: The spectrum gave a peak at 370 0nm, confirming the formation of zinc oxide nanoparticles

**Figure 3.7.**
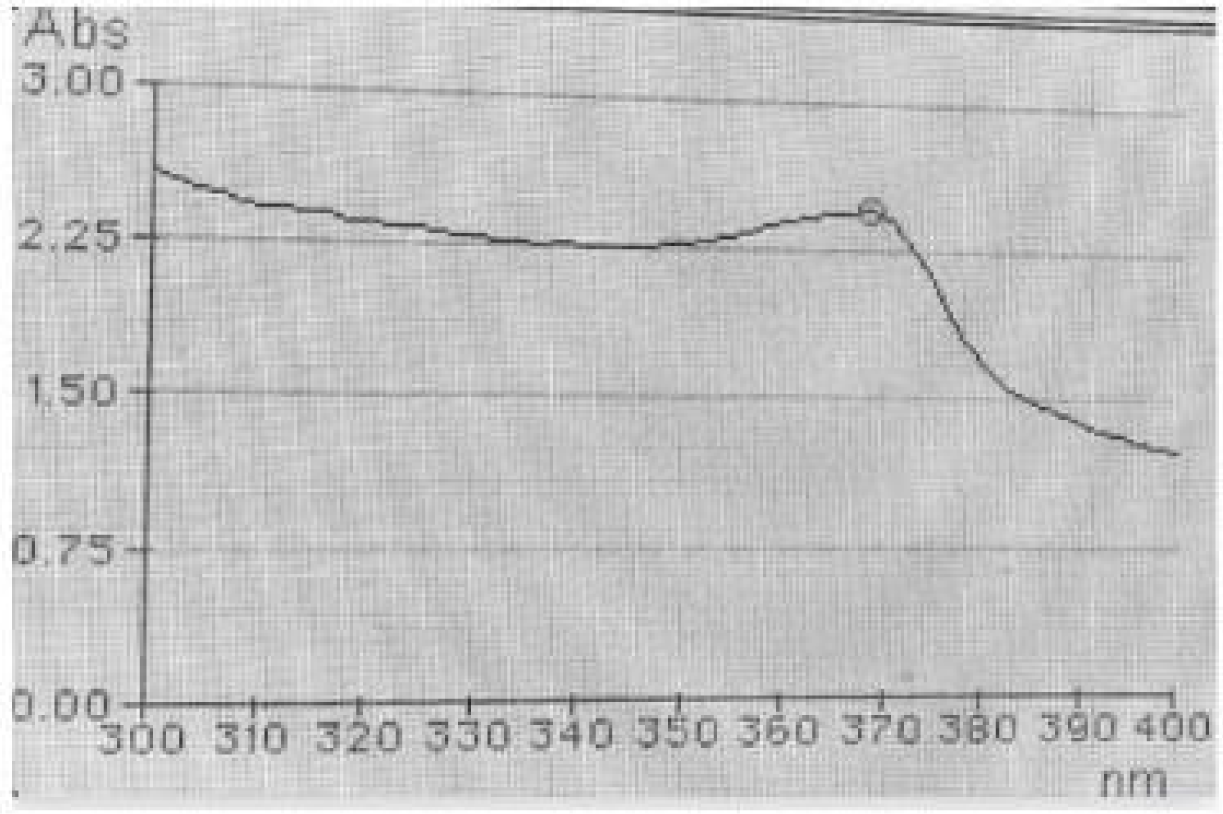
UV/Vis Spectroscopy results of cobalt-doped Zinc Oxide nanoparticles: **The** :spectrum gave a peak **at** 368nm, comrunmg the formation of cobalt-doped zinc oxide nanoparticles

**Figure 3.8.**
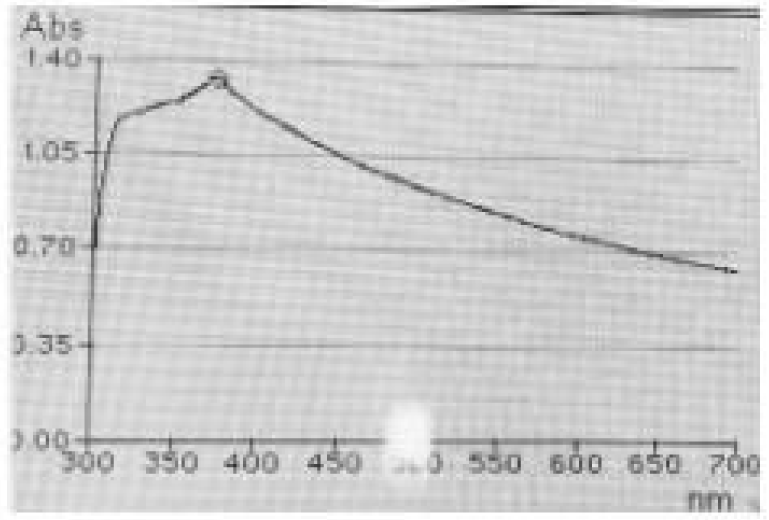
UV/Vis Spectrosrceosupltys of Manganese doped Zinc Oxide nanoparticles: The spectrum gave a peak at 373nm, confirming the formation of Manganese doped zinc oxide nanoparticles

**Figure 3.9.**
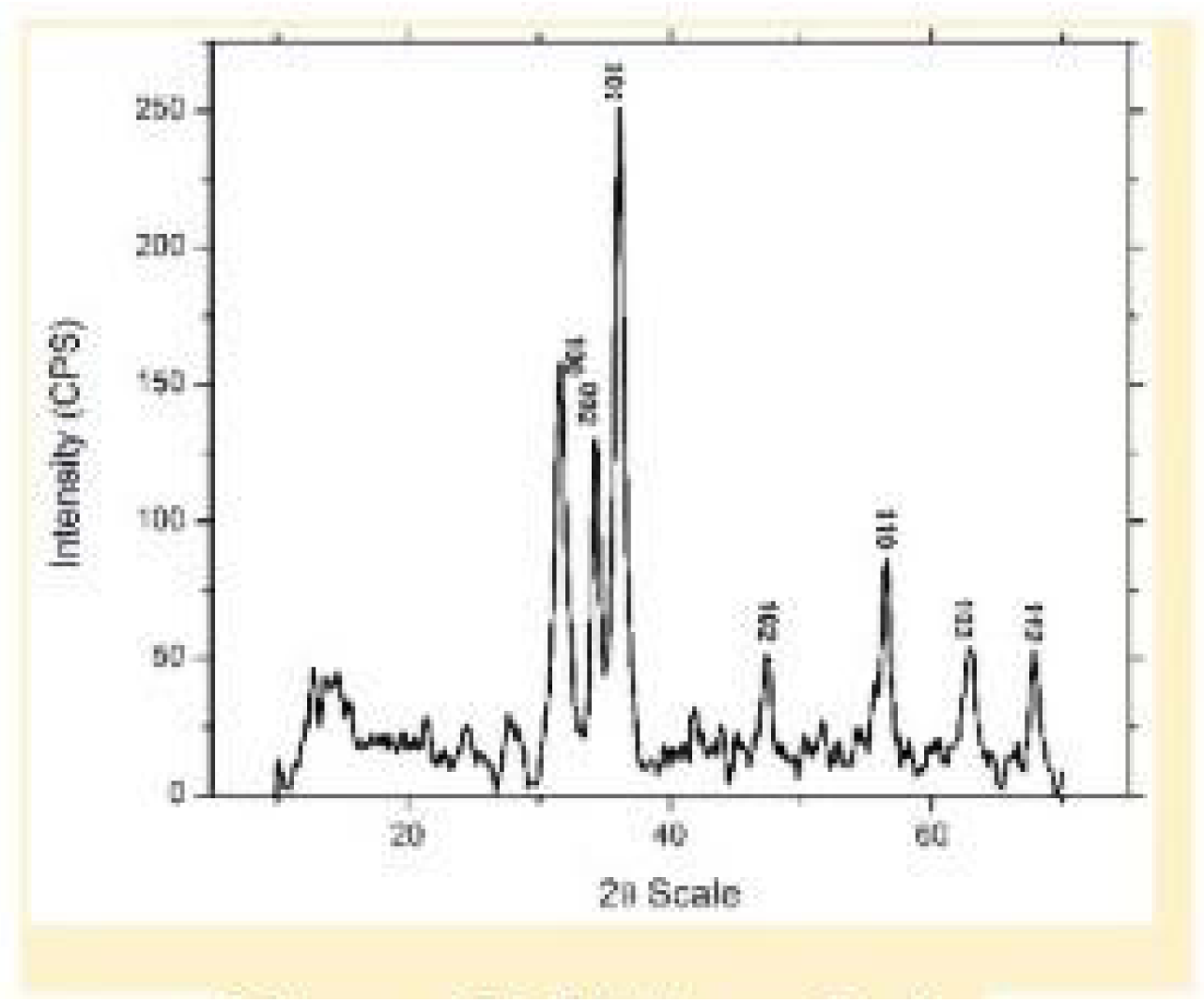
XRD analysis

**Figure 3.10.**
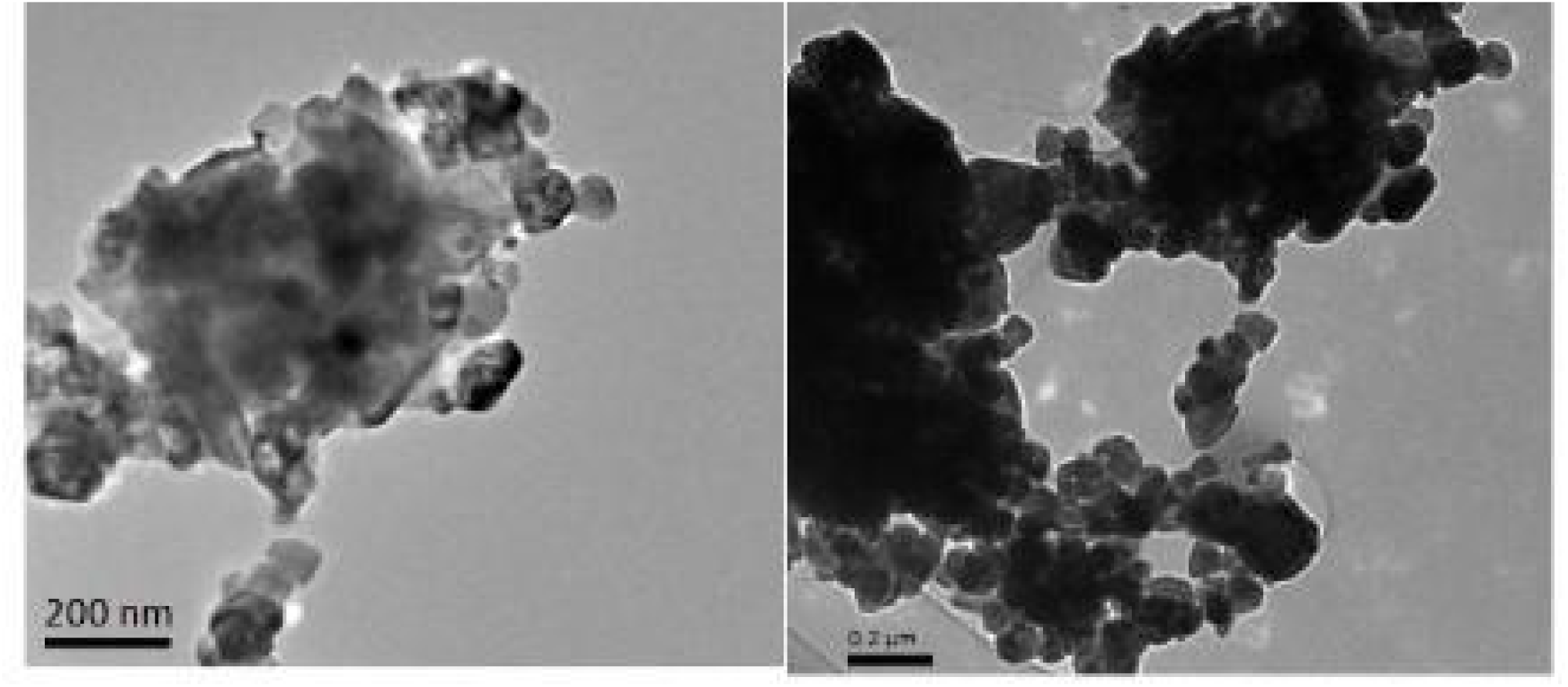
Transmission electron microscope result

**Figure 3.11.**
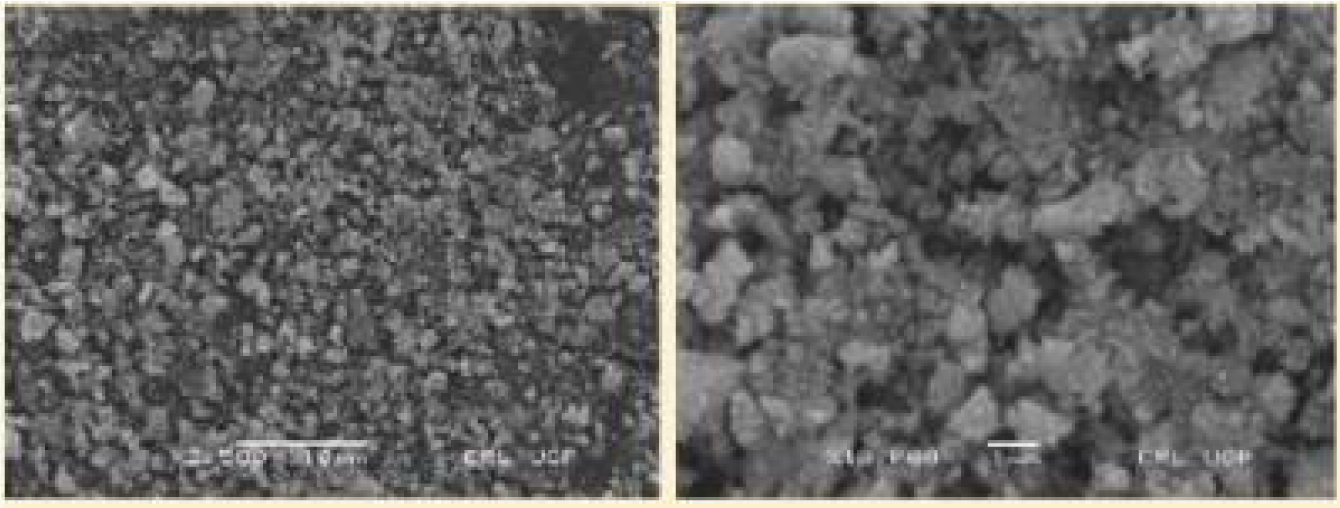
SEM analysis

**Figure 3.12.**
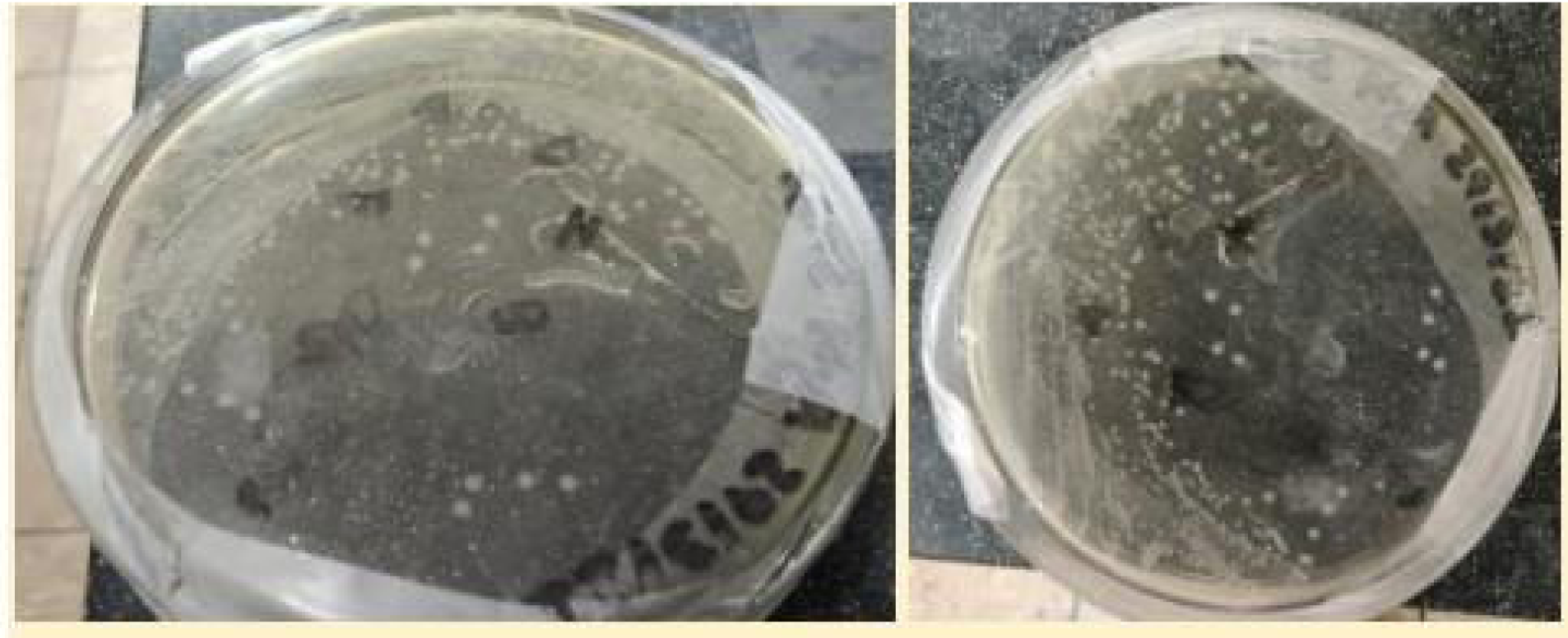
Streak Plate Method Image (Isolated Colonies of *E. coli*)

**Figure 3.13.**
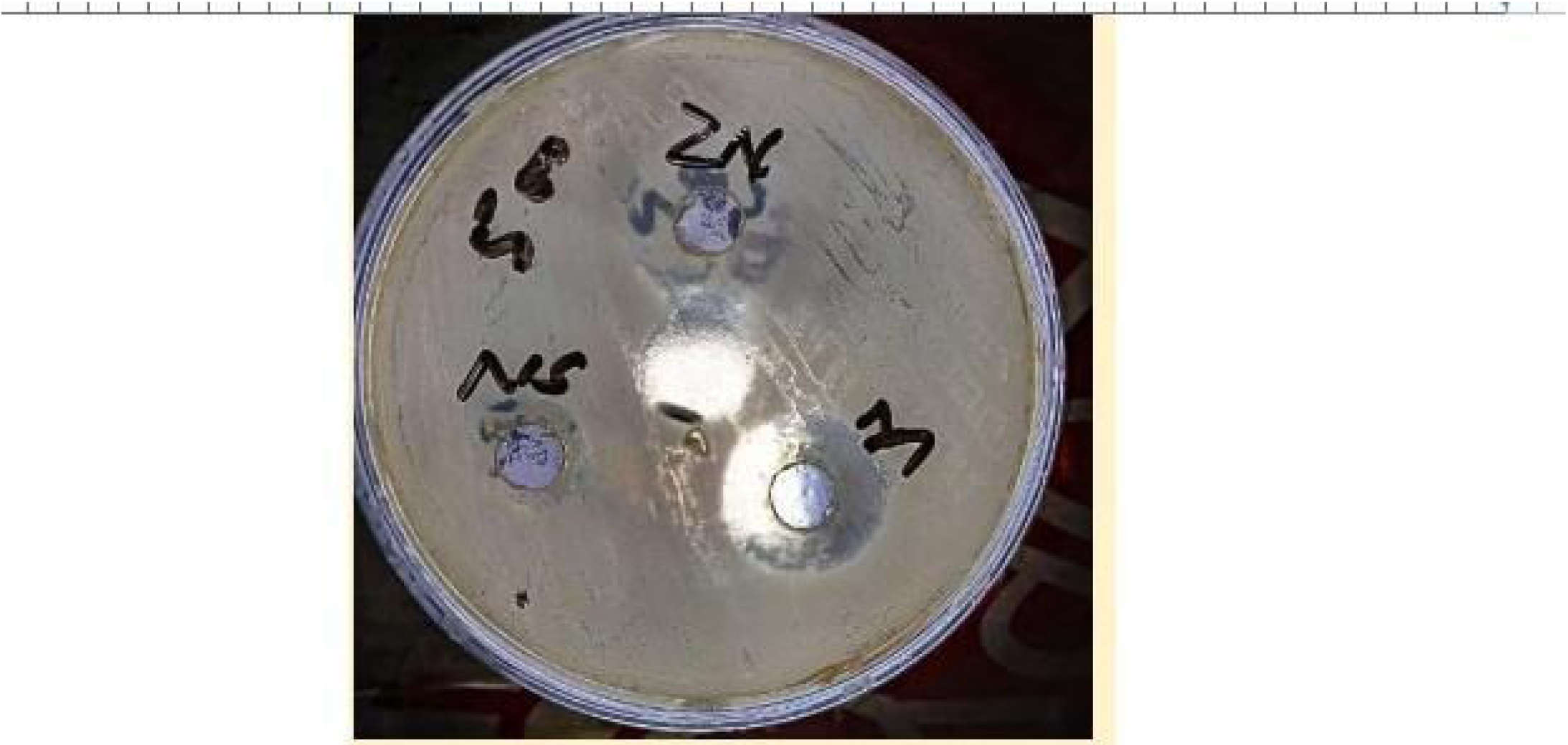
Agar well diffusion method of zinc oxide nanoparticles against *E. coli* Inhibition zone was: 2.5cm(25mm) on zinc oxide, 1.3(13mm) on *Nigella* sativa and 2.6cm(26mm) on combine effect

**Figure 3.14.**
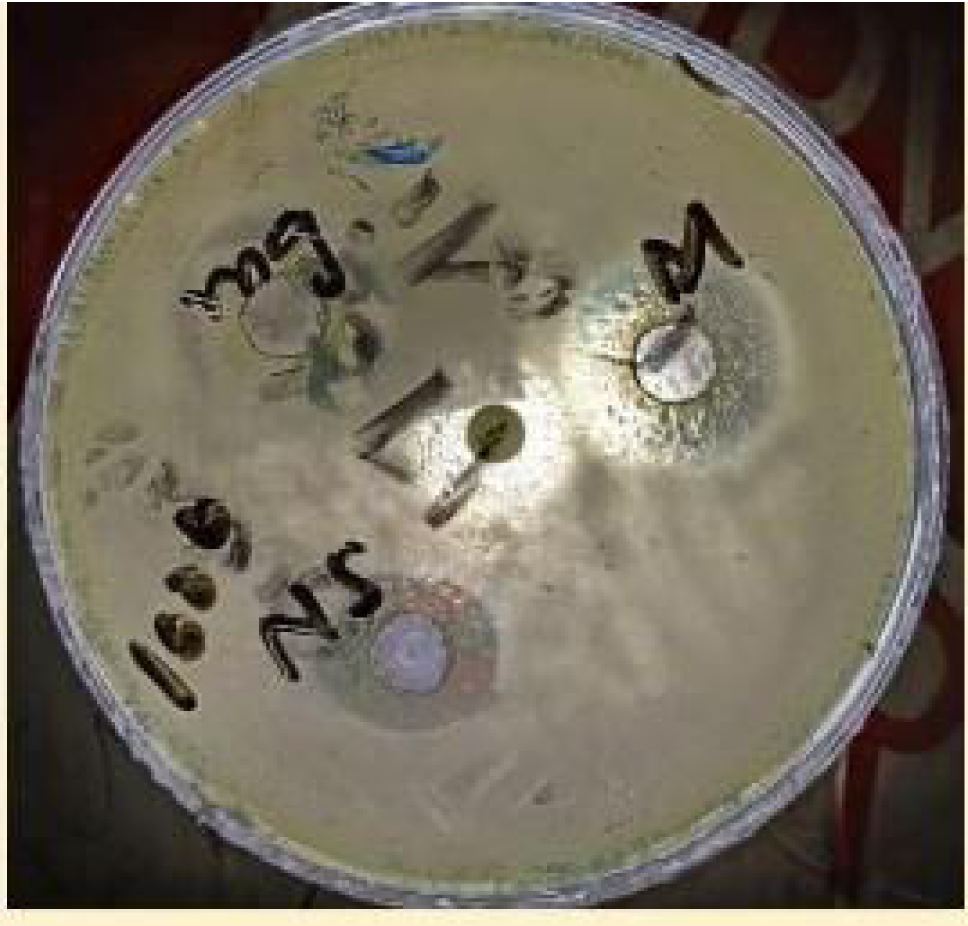
Agar well diffusion method of manganese doped zinc oxide nanoparticles against *E. coli*

**Figure 3.15.**
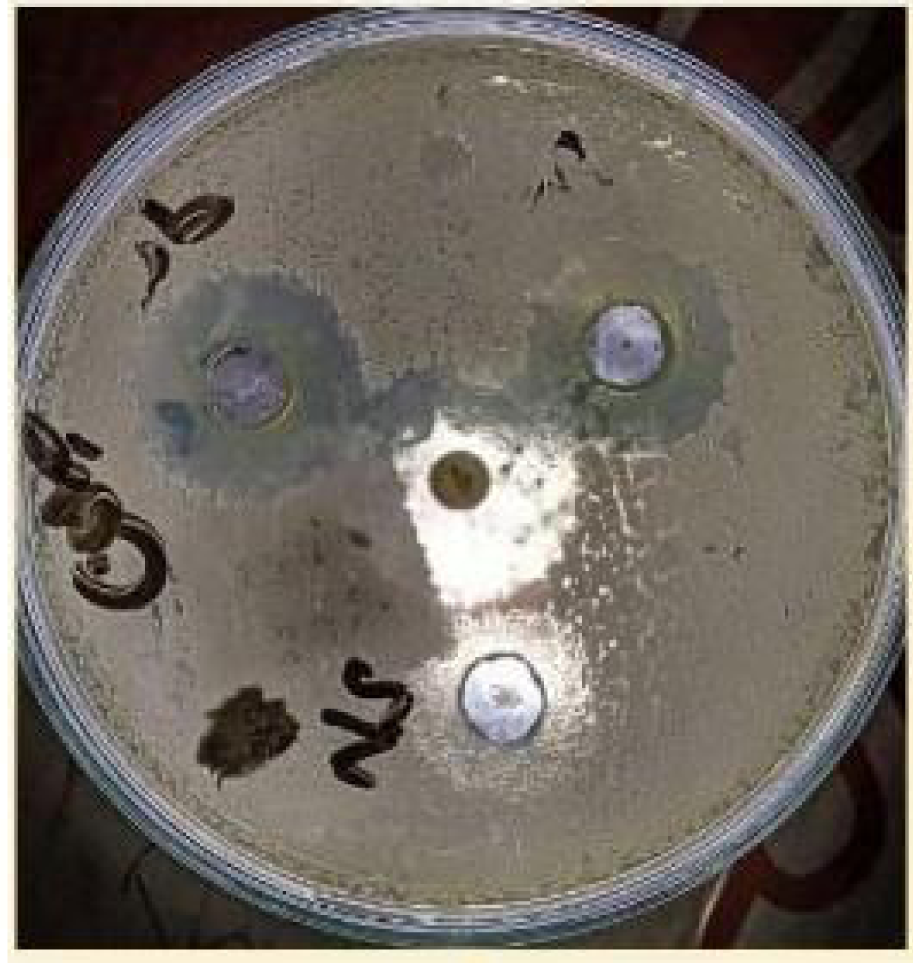
Agar well diffusion method of cobalt doped zinc oxide nanoparticles against *E. coli*

**Figure 3.16.**
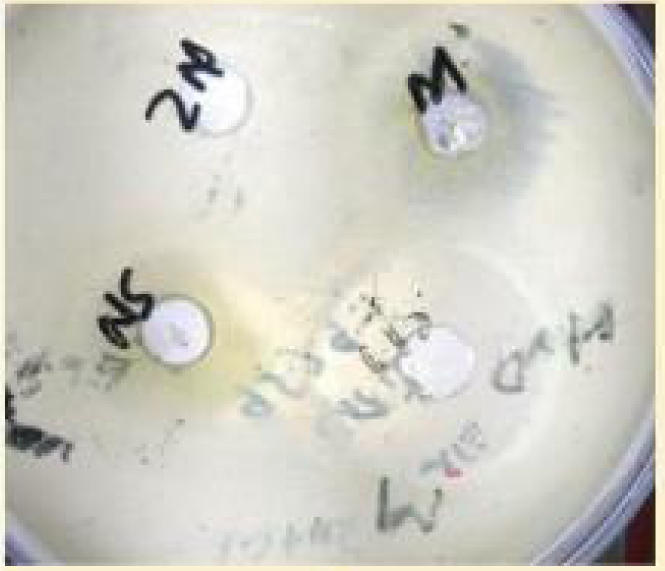
Agar well diffusion method of Ethanol against *E. coli* The inhibition zone was: 3.0cm(30mm)

**Figure 3.17.**
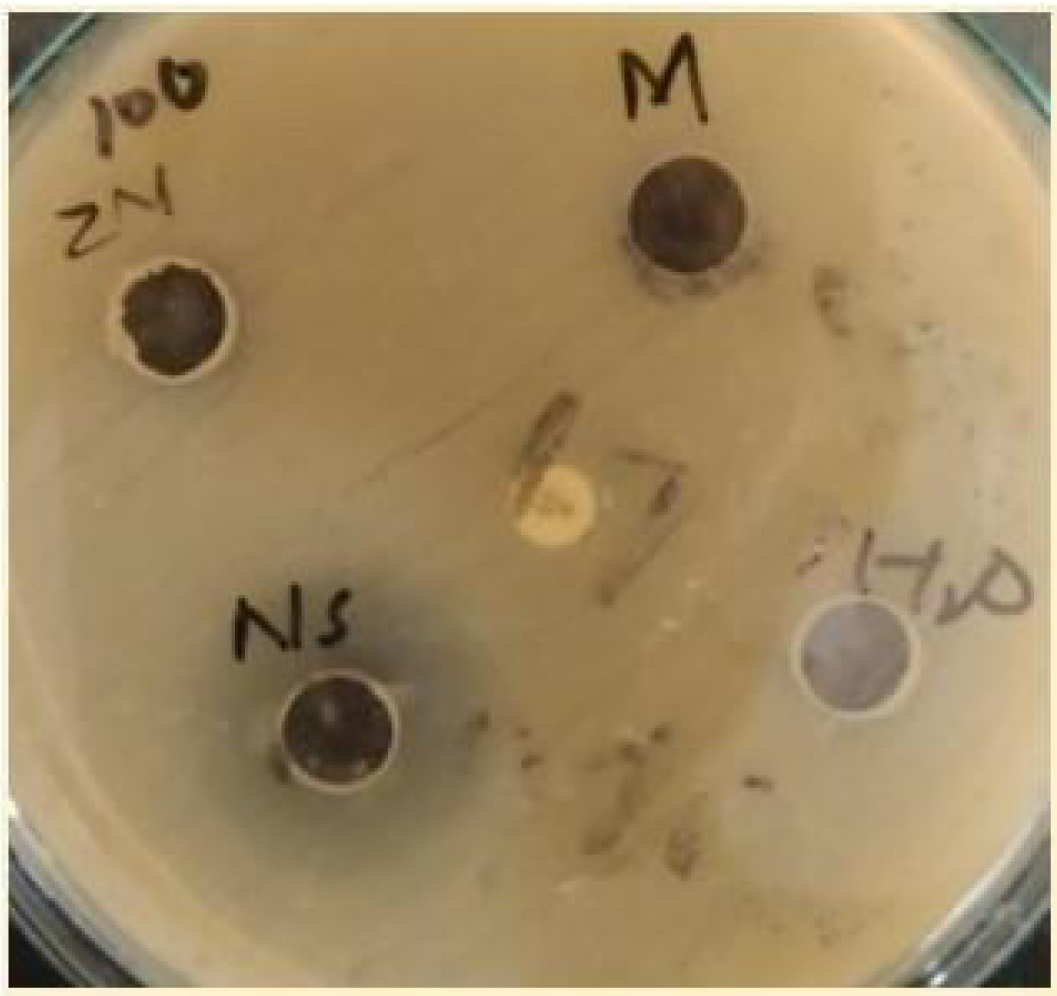
Agar well diffusion method of zinc oxide nanoparticles against Staph. Aureus

**Figure 3.18.**
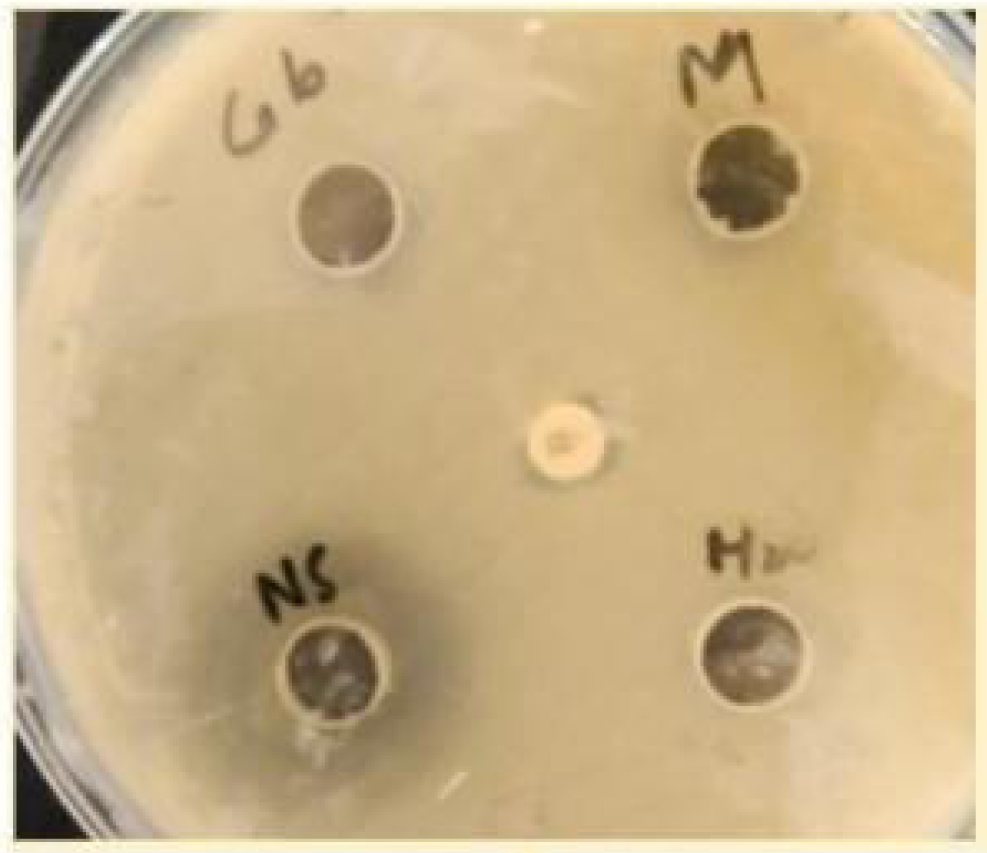
Agar well diffusion method cobalt doped zinc oxide nanoparticles against *Staph aureus* The inhibizotnei woasn: 0 cm on zinc oxide, 2.0cm(20mm) on *Nigella* sativa, and 0cm on the combined effect The inhibition zone was: 0 cm on zinc oxide, 2.0cm(20mm) on Nigella sativa, and 0cm on the combined effect

**Figure 3.19.**
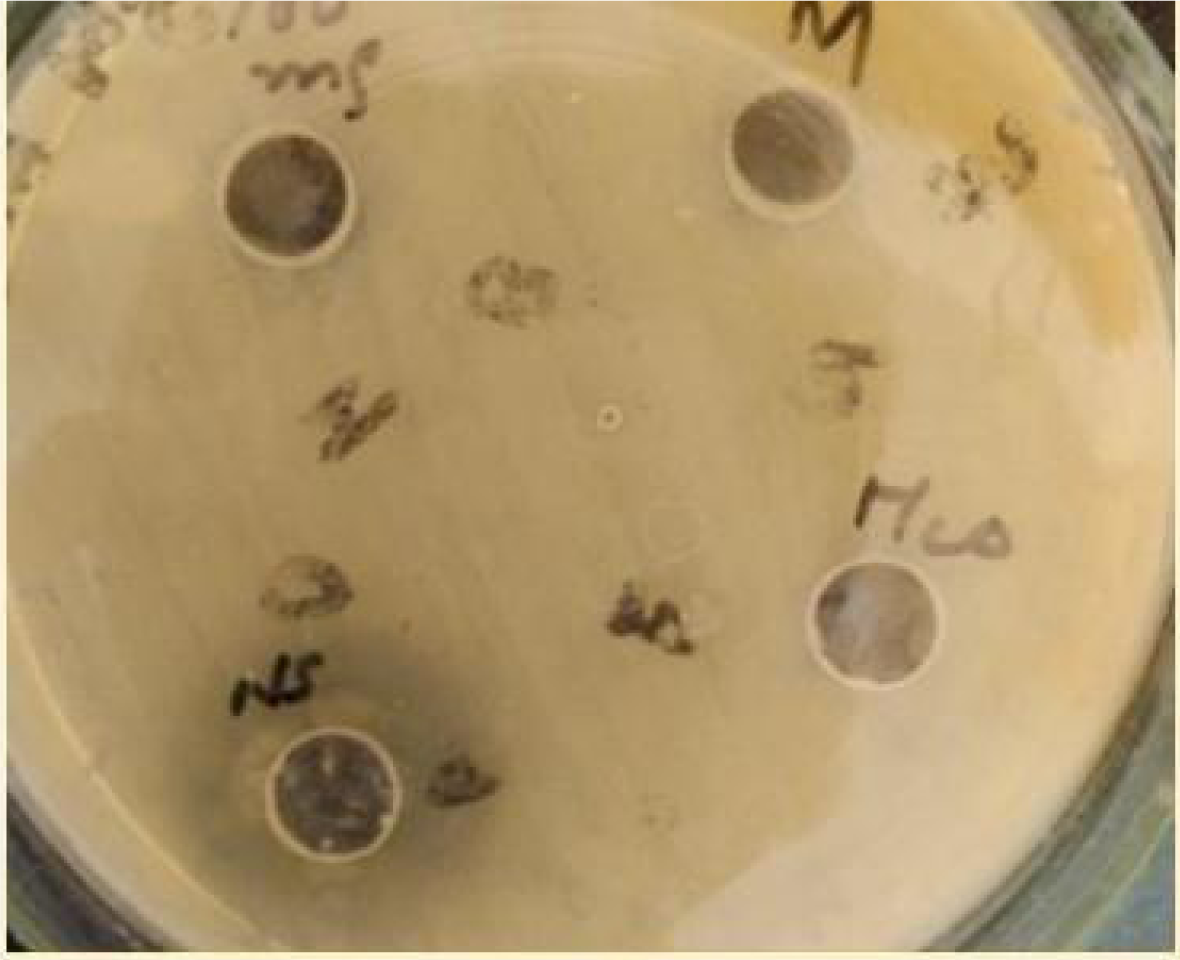
Agar well diffusion method of manganese doped zinc oxide nanoparticles against Staph aureus The inhibition zone was: 0 cm on zinc oxide, 2.0cm(20mm) on Nigella sativa, and Ocm on the combined effect

**Figure 3.20.**
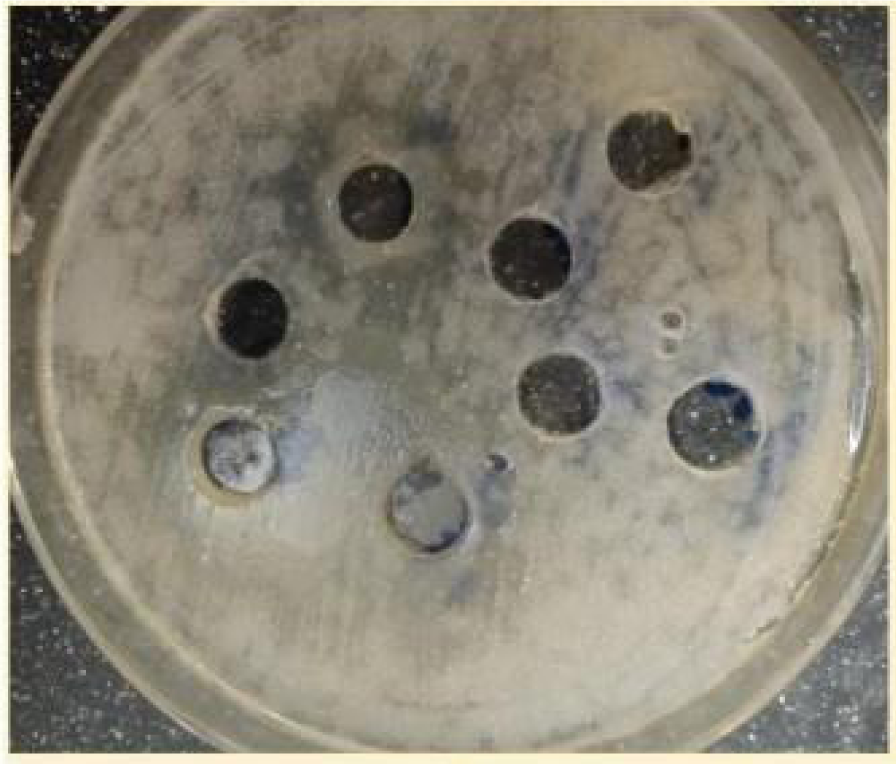
synergetic effects of zinc oxide and *Nigella sativa*= 2.1cm inhibition zone cobalt doped zinc oxide and Nigella sativa=2.2cm inhibition zone manganese doped zinc oxide and *Nigella sativa*=2.1cm inhibition zone Bacteria= Staph aureus

**Figure 3.21.**
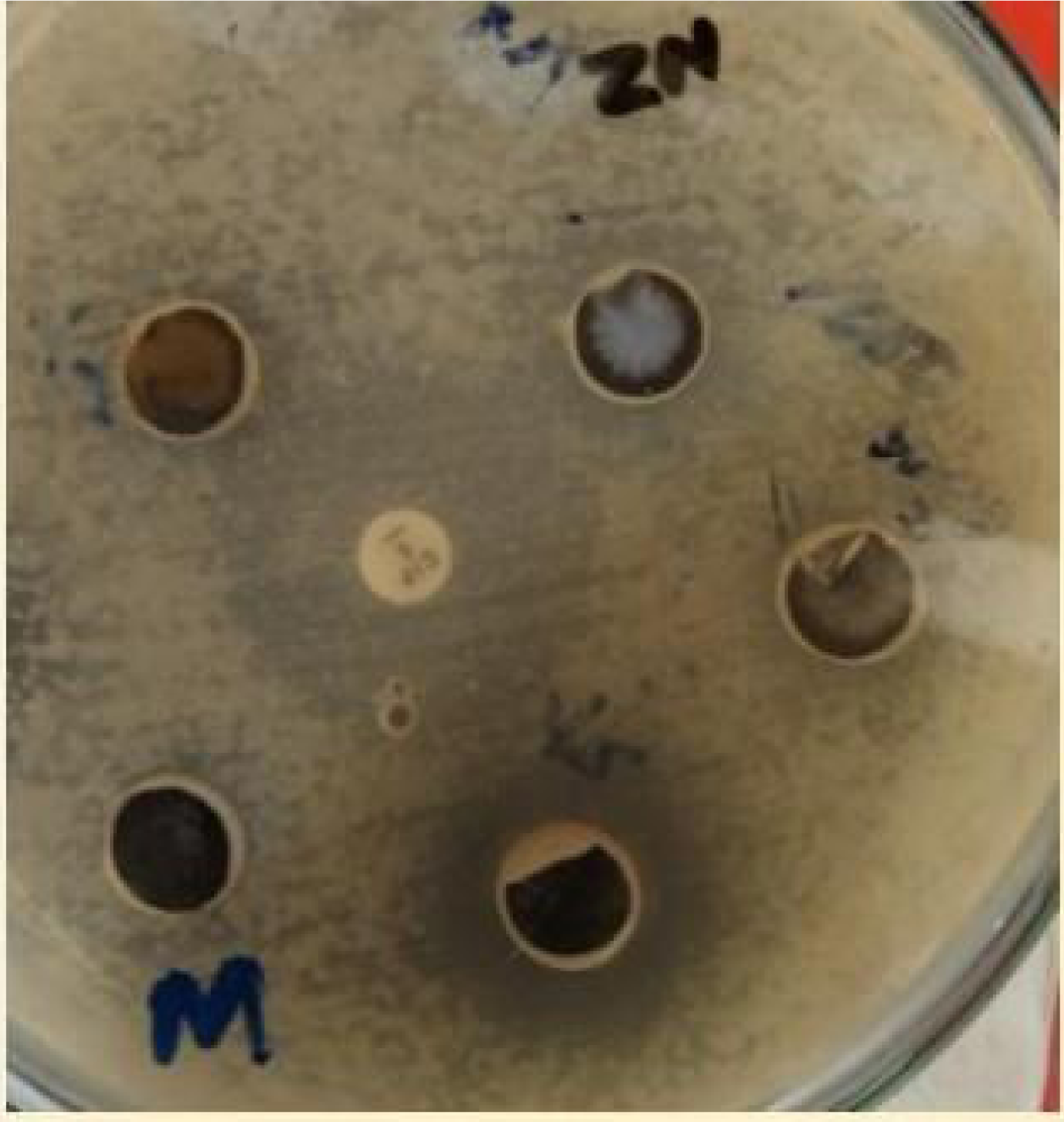
*Nigella sativa* effect against *Pseudomonas* Inhibition zone was = 2.2cm(22mm) No synergetic effect No NPs effects were seen

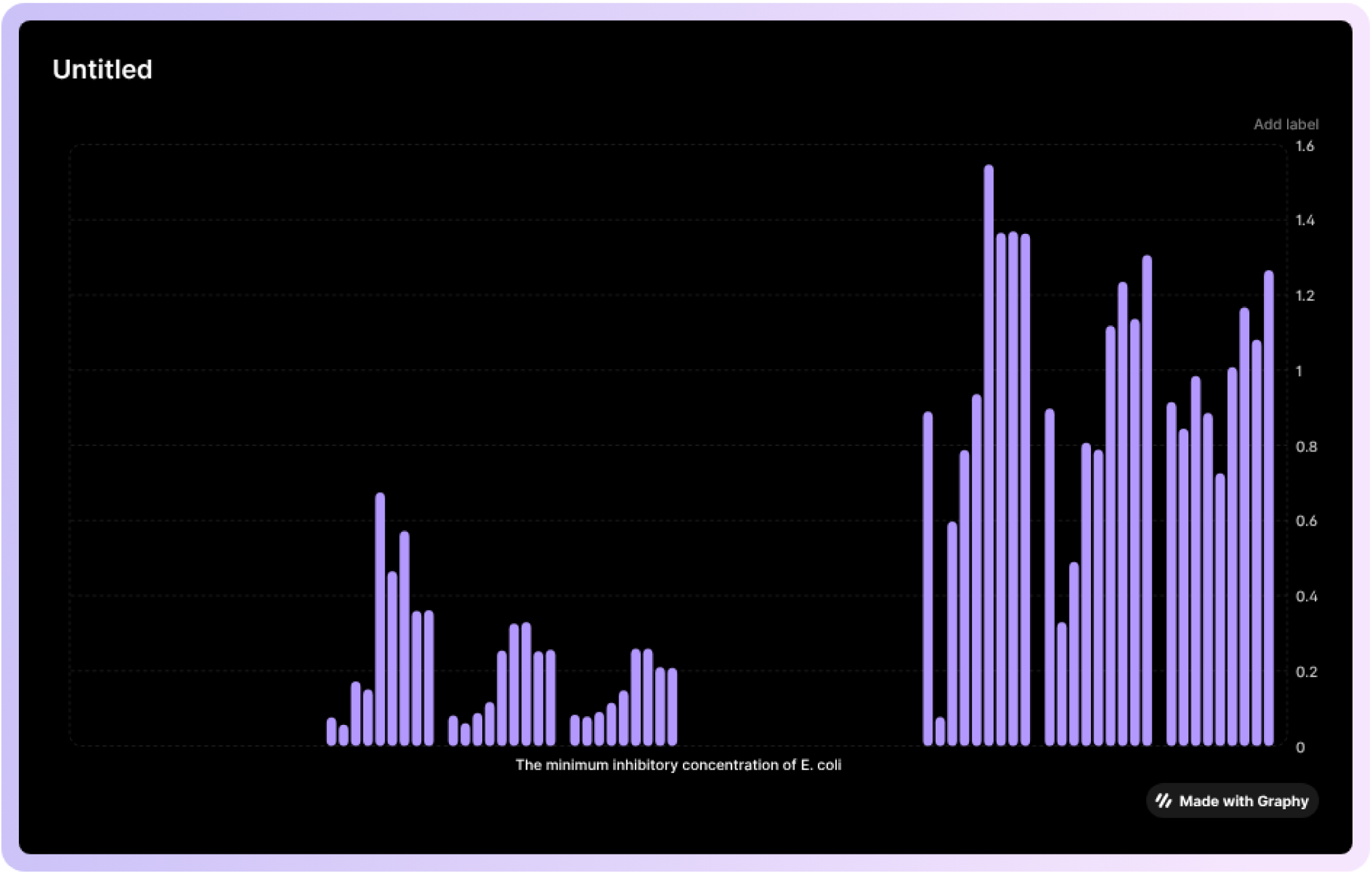

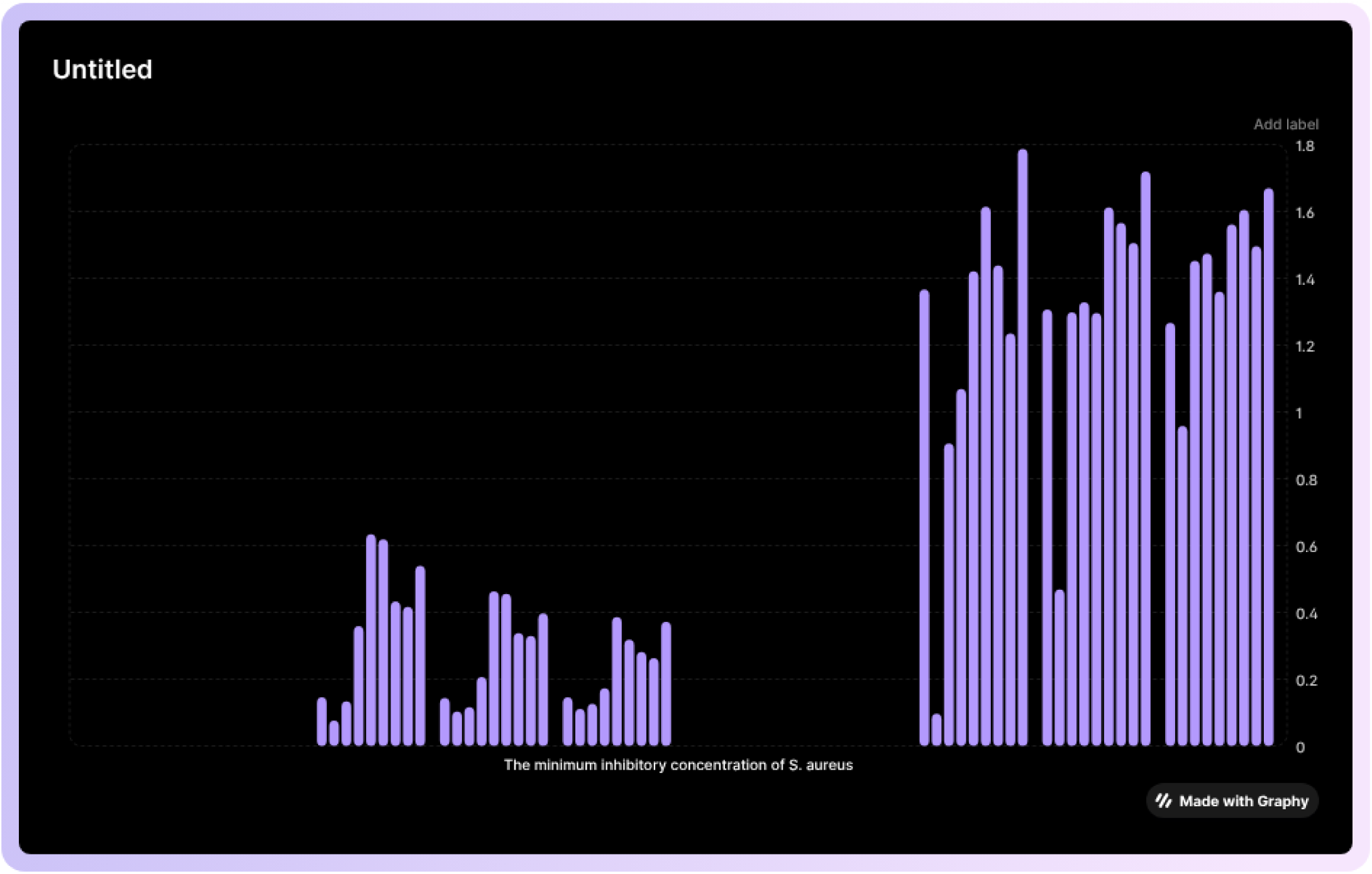

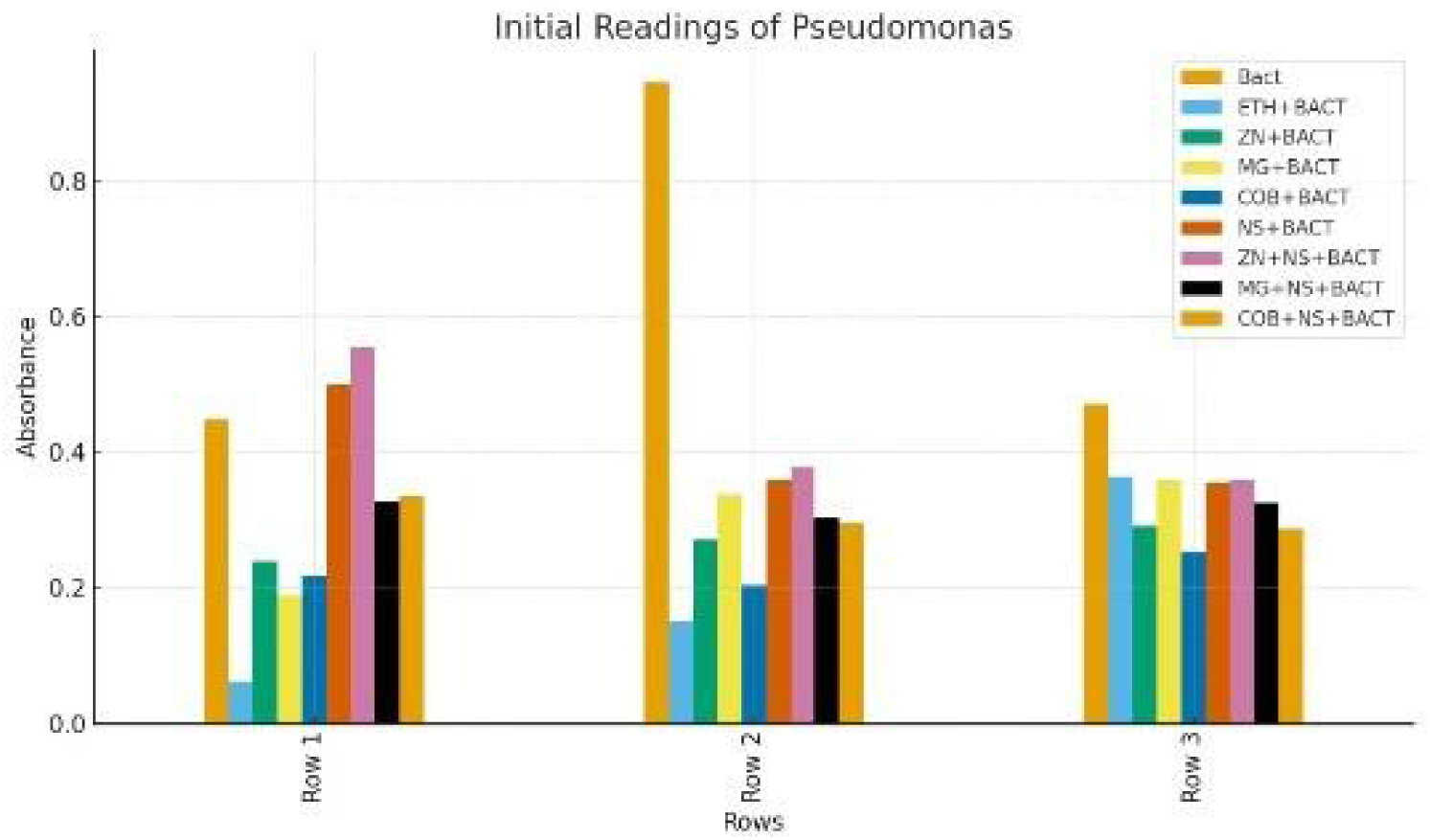

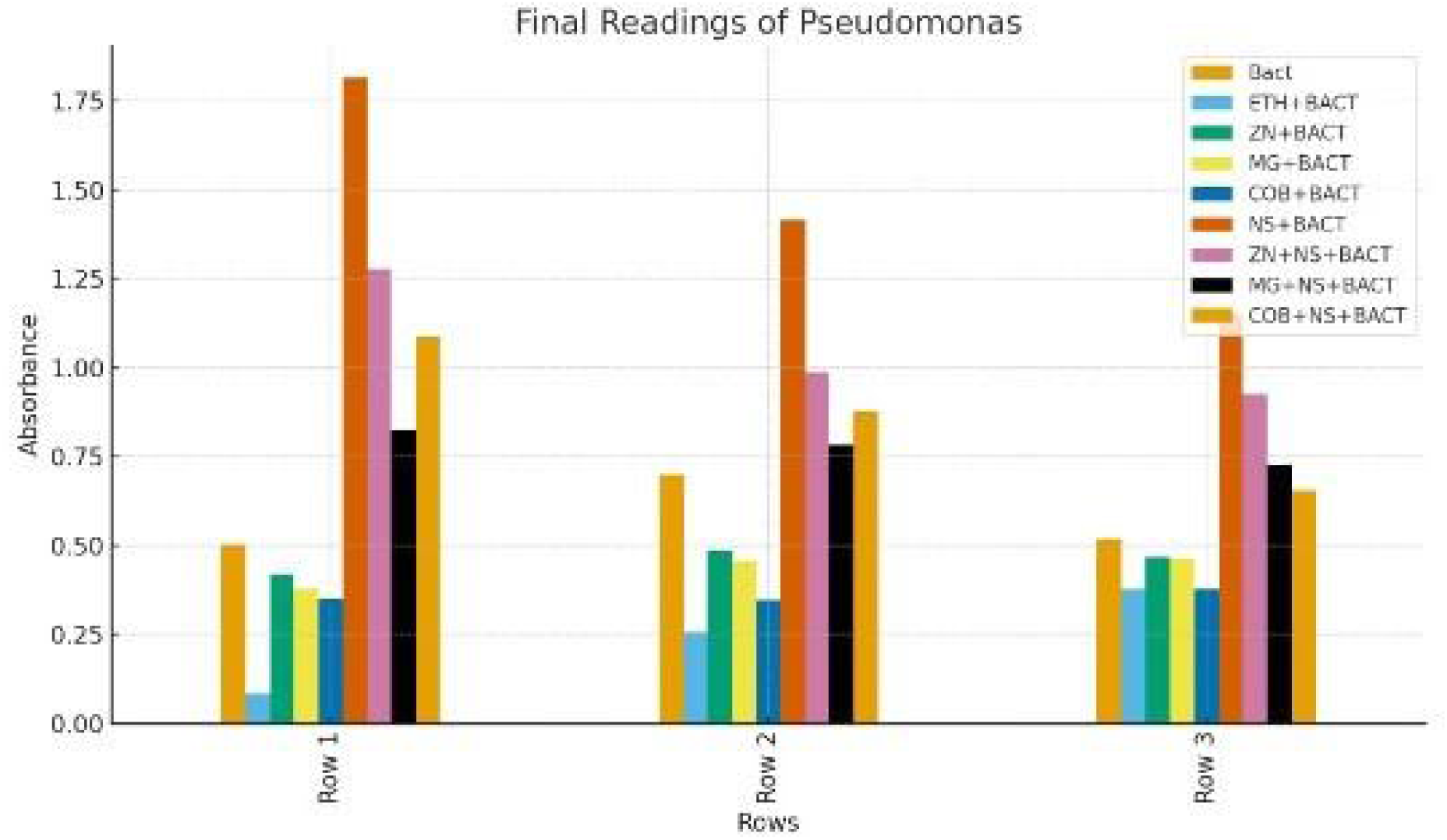

